# Machine learning reveals missing edges and putative interaction mechanisms in microbial ecosystem networks

**DOI:** 10.1101/286641

**Authors:** Demetrius DiMucci, Mark Kon, Daniel Segrè

**Author notes:** Correspondence to Daniel Segrè.

## Abstract

Microbes affect each other’s growth in multiple, often elusive ways. The ensuing interdependencies form complex networks, believed to influence taxonomic composition, as well as community-level functional properties and dynamics. Elucidation of these networks is often pursued by measuring pairwise interaction in co-culture experiments. However, combinatorial complexity precludes the exhaustive experimental analysis of pairwise interactions even for moderately sized microbial communities. Here, we use a machine-learning random forest approach to address this challenge. In particular, we show how partial knowledge of a microbial interaction network, combined with trait-level representations of individual microbial species, can provide accurate inference of missing edges in the network and putative mechanisms underlying interactions. We applied our algorithm to two case studies: an experimentally mapped network of interactions between auxotrophic *E. coli* strains, and a large *in silico* network of metabolic interdependencies between 100 human gut-associated bacteria. For this last case, 5% of the network is enough to predict the remaining 95% with 80% accuracy, and mechanistic hypotheses produced by the algorithm accurately reflect known metabolic exchanges. Our approach, broadly applicable to any microbial or other ecological network, can drive the discovery of new interactions and new molecular mechanisms, both for therapeutic interventions involving natural communities and for the rational design of synthetic consortia.

**Importance:** Different organisms in a microbial community may drastically affect each other’s growth phenotype, significantly affecting the community dynamics, with important implications for human and environmental health. Novel culturing methods and decreasing costs of sequencing will gradually enable high-throughput measurements of pairwise interactions in systematic co-culturing studies. However, a thorough characterization of all interactions that occur within a microbial community is greatly limited both by the combinatorial complexity of possible assortments, and by the limited biological insight that interaction measurements typically provide without laborious specific follow-ups. Here we show how a simple and flexible formal representation of microbial pairs can be used for classification of interactions with machine learning. The approach we propose predicts with high accuracy the outcome of yet to be performed experiments, and generates testable hypotheses about the mechanisms of specific interactions.

## Introduction

The collective behavior of microbial ecosystems across biomes is an outcome of the many interactions between members of the community (1, 2). These interactions include exchange of metabolites, signaling and quorum sensing processes, as well as growth inhibition and killing. Understanding the interspecific interactions within microbial communities is essential for understanding the function of natural communities (3) and for the design of synthetic communities (4).

A powerful, and increasingly employed method for assessing intermicrobial interactions is the direct measurement of phenotypes of microbial species grown in co-culture (5–7). A fundamental challenge in this endeavor is the huge diversity of many natural communities, which could count up to several hundred strains or species of microbes. Performing experiments for all possible pairwise interactions constitutes a herculean, and likely insurmountable task for even a moderately sized community. It is however, conceivable that new computational approaches could systematically complement existing tools such as high-throughput sequencing and genome annotation (8–13) to help extract as much information as possible from interaction datasets, providing both insight on yet-to-be-measured interactions, and on possible biological mechanisms mediating specific partnerships.

Here we present a conceptual framework for the mathematical representation of microbial interactions and subsequent use of supervised learning to build a classifier with high predictive accuracy. While any algorithm may be used, we obtained our best results with random forest (14). Random forests are ensembles of many decision trees that individually are poor classifiers but can be democratically pooled to create a very good classifier. Random forests have two attributes that we found particularly attractive for our purposes here. First, they are non-parametric and thus require no *a priori* definition about underlying relationships between predictive variables. Second, recent methodological developments in the interpretation of random forests have been made that allow users to query why a specific example was classified the way that it was through the calculation of feature contributions (15). Feature contributions can be exploited to develop new hypotheses about the mechanisms that mediate specific interactions. In order to demonstrate a proof of principle for the classification of microbial interactions using organism traits and the utility of feature contributions for developing insight into the underlying mechanisms, we applied this approach to two communities where all pairwise experiments had been performed and the mechanisms of interaction identified. The first is an *in silico* community of 100 metabolic models of human gut associated bacteria and the second are the experimental results of a study involving 14 amino acid auxotrophic strains of *Escherichia coli* (*E. coli*). Our results show that the application of random forests and feature contributions to the study of microbial communities can significantly increase the capacity to estimate a large number of interactions based on a limited number of experiments, and has the potential to enable the discovery of new interaction mechanisms.

## Results

### Representing Pairwise Interactions

Our objective in this study was twofold; first, we sought to predict the qualitative outcome of unobserved pairwise interactions in a microbial community; second, we wanted to identify predictive variables that suggest potential mechanisms of interaction. In order to achieve both of these goals it was important to establish a representation that enables an algorithm to make good predictions and also be easily parsed for interpretation. Our approach relies of the availability of a trait-level description available for each organism in the community under consideration. Trait descriptions are used to construct trait vectors for each organism. Specific interactions are represented as the concatenation of the relevant trait vectors (Figure 1). Trait vectors may be constructed from any set of biologically relevant features such as body size or metabolic requirements; here we used genome-scale metabolic data for every organism in the *in silico* community case study, and vectors of biosynthetic capabilities for each *E. coli* strain in the auxotroph community case study.

**Figure 1.**
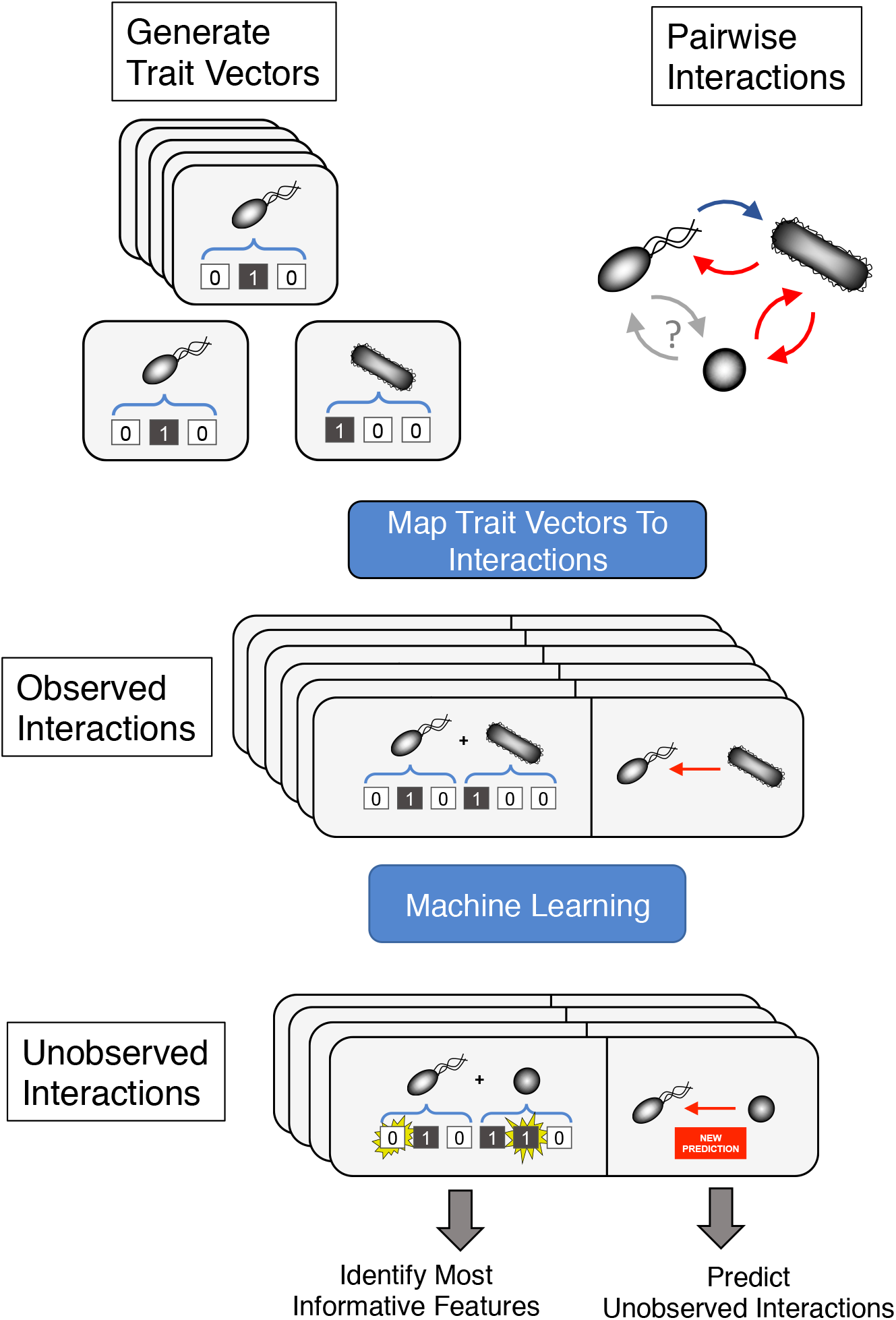
A schematic representation of our machine learning approach for inferring interactions among microbes. A trait vector captures the characteristics of each organism in the community of interest. The presence or absence of a trait in a given organism is encoded (as a binary number) in the corresponding element of the trait vector. For every possible pairwise interaction among community members we construct a composite vector that is the concatenation of the corresponding trait vectors. The vector of the organism whose response is being predicted is concatenated to the front of the trait vector of its interaction partner. For the set of observed interactions each composite vector is then mapped to the measured response of the interacting species. All observed interactions are then used to train a model that predicts the outcome of unobserved interactions. If random forest is used then feature contributions can be calculated on a case-by-case basis in order to identify which elements of the composite genome contribute most strongly to the prediction.

These trait vectors, together with the known outcome of specific interactions, can be fed into a machine learning algorithm capable of absorbing existing patterns and extending predictions to unknown interactions. The approach we use here, a random forests classifier, is an ensemble of many decision trees that individually ask a series of yes or no questions about randomly selected subsets of predictive features in order to classify samples. Single trees tend to be poor classifiers; however, when their decisions are pooled, the collective predictive power is often impressive. In order to find potential mechanisms of interaction we took advantage of the binary nature of individual trees in order to identify which variables are the most influential for the classification of specific samples.

### Application to computationally predicted interactions between human gut microbes

We first applied our approach to a large *in silico* data set that we generated by simulating with dynamic flux balance analysis (16) (using COMETS (17), see Methods) all pairwise co-culture interactions between 100 metabolic models of human-gut associated bacteria (18) under rich medium, in a well-mixed batch culture. Each metabolic model is a network of over a thousand interconnected metabolic reactions that fall into two categories: intracellular and exchange reactions (19). Intracellular reactions correspond to the various pathways found within the cell, while exchange reactions represent the ability of an organism to transport a metabolite across the cell membrane. Among the 100 organisms, there were a total of 2083 unique metabolic reactions that could serve as potential predictors. Out of these, 194 were exchange reactions. Since interactions between organisms emerge through the exchange of or competition for one or more extracellular metabolites in the environment, we used only the exchange reactions as predictive features (Figure 2A).

**Figure 2.**
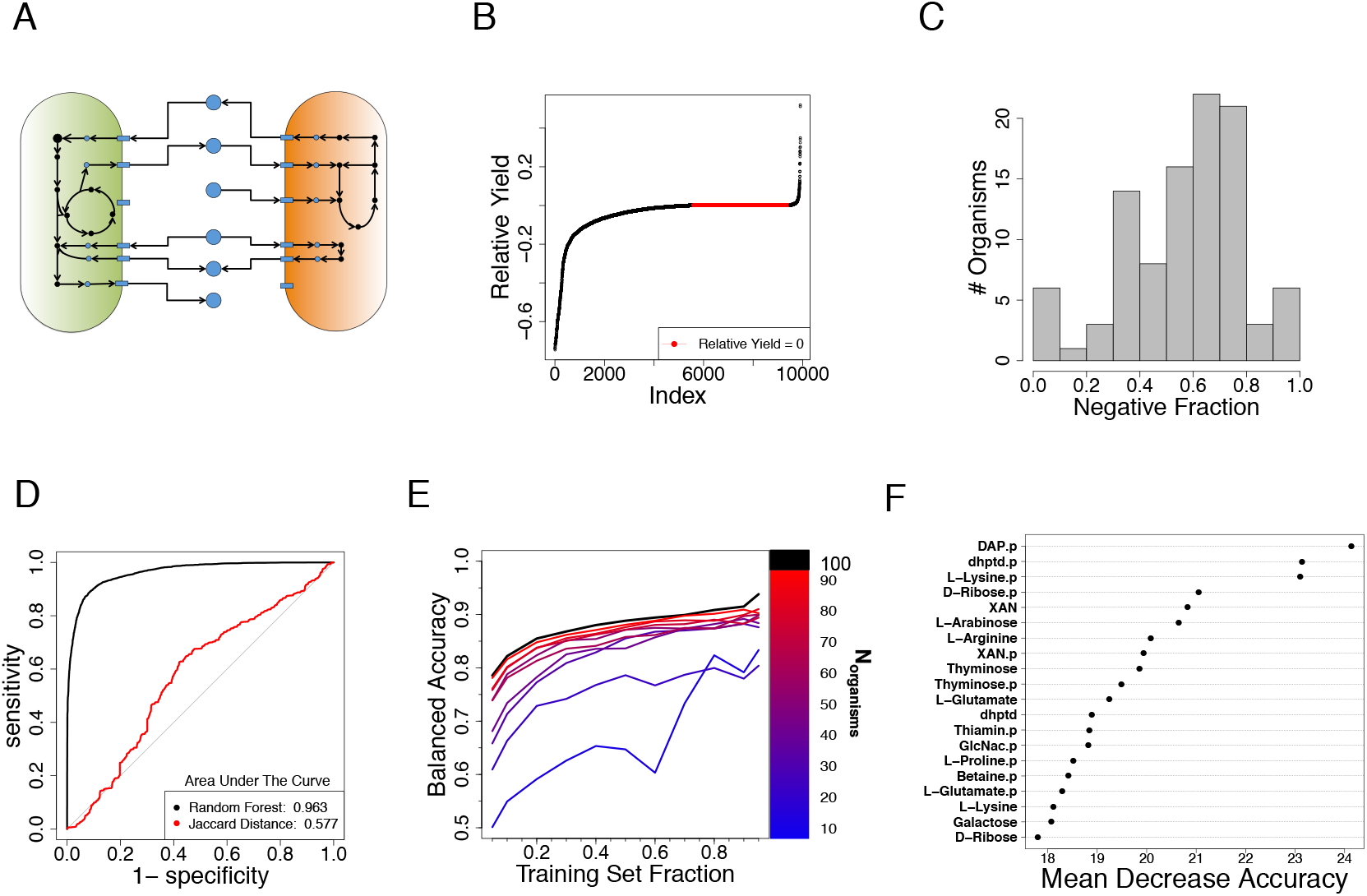
Classification of interactions in an *in silico* model of a community of human gut microbes. **A.** Organisms are represented *in silico* as large networks of metabolic reactions that take up metabolites (blue circles) from the environment (arrows leading to model) and release by-products (arrows leading to metabolite). Organisms may interact with one another during simulation when both organisms compete for the uptake of a metabolite or through cross feeding where one model consumes a by-product of the other. **B.** The distribution of relative yields is sorted and plotted. Interactions resulted in a model producing a negative relative yield 5563 times. Neutral interactions, a relative yield of zero, occurred 3917 times, and positive relative yield happened 420 times. Samples were classified as negative or non-negative. **C.** Histogram of the fraction of interactions that were the negative class for each organism. The distribution is consistent with a truncated normal distribution centered on .562 (t test, p ≈ .67) **D.** ROC curve of a random forest classifier using 388 exchange reactions as predictors for 9900 *in silico* observations compared to the ROC curve obtained from using Jaccard Distance as a threshold to predict negative versus non-negative relative yield. **E.** Learning curves for sub-communities of the full *in silico* community. These learning curves are the median learning curve for 5 sub-communities selected at random for each value of Norganisms. **F.** 20 most influential predictors as determined by mean decrease in accuracy. The ‘.p’ suffix indicates that the predictor belongs to the interaction partner. Some metabolite names are shortened: DAP = meso-2,6-Diaminopimelate, dhptd = 4-5-dihydroxy-2-3-pentanedione, XAN = xanthine, GlcNac = N-Acetylglucosamine.

Interactions in the network were computed by determining the influence of each organism on each other organism in co-culture simulations. We compared the accumulated biomass of each model in co-culture experiment to its accumulated biomass in monoculture and expressed the phenotypic response as relative yield (Methods). A negative relative yield indicates that a model produces less biomass in co-culture than it does when growing alone, a relative yield of 0 indicates that its growth is unaffected by interaction with another model, and a positive relative yield means that the model benefits from the presence of another. For the purpose of binary classification, we divided observations into two classes based on their relative yields. Samples with a negative relative yield were classified as ‘negative’ responders (5563/9900). Samples with a positive relative yield (420/9900) or a relative yield of zero (3917/9900) were classified as ‘nonnegative’ responders (Figure 2B). We refer to the model whose response is being predicted as the ‘responder’ and its interaction partner is the ‘partner’. In order to develop an intuition for how difficult this classification task might be we calculated what fraction of the 99 interactions each responder took part resulted in a negative relative yield. Classification could become a trivial task if the distribution of these fractions were bimodal with peaks far from each other, the most extreme scenario being one where responders experience either only negative or non-negative relative yields in co-culture. On the other hand, a normal or uniform distribution would indicate a more complicated reality where classification would benefit from machine learning. This last option was indeed confirmed to hold for our data, as the histogram of the negative fractions follows a truncated normal distribution (Figure 2C).

In order to establish a baseline for comparing the effectiveness of the random forest as a classifier, we calculated the Jaccard distance of each interaction pair and then used the Jaccard distance as a decision threshold for predicting whether a model would have a negative or non-negative relative yield in co-culture. Jaccard distance was chosen as a baseline predictor because it is easy to calculate and is commonly used in statistical analyses of microbial communities to infer interactions (20, 21). The resulting receiver operator characteristic (ROC) curve was then compared to the ROC curve obtained from a random forest classifier trained using the 194 exchange reactions as features to predict the outcome the full set of 9900 observations. As a classifier, the Jaccard distance does not perform much better than randomly guessing, while the random forest was surprisingly good (Figure 2D). To evaluate the accuracy of the random forest classifier on the full data set we examined the out of bag (OOB) error estimates. OOB estimates in random forests have been shown to be roughly equivalent to five-fold cross-validation (22). We therefore took the OOB error rate as an estimate for the overall test error rate. The random forest resulted in a balanced accuracy of 90.4%.

High predictive accuracies are encouraging but will be of little use if they can only be achieved when the vast majority of the experiment outcomes are already known. Learning curves visualize how the performance of a classifier behaves as the data available for trainings increases. We constructed a series of learning curves to measure how the balanced accuracy of the random forest classifier is affected by the size of the community being studied and as the fraction of interaction outcomes known increases (Figure 2E). As is typically seen with learning curves, classifier performance improves when more of the data is made available. Interestingly we observe that for the smallest communities (*N_organisms_* = 10) balanced accuracy was better than random when as little as 10% of the data was available (2 experiments), but the trajectory with additional data improves at a much slower rate than it does for larger communities. When *N_organisms_* is increased to 20, significant improvements to balanced accuracy manifest immediately and have a stronger benefit with the outcome of additional experiments. Encouragingly, these benefits appear moderate when compared to the trajectory of much larger communities (*N_organisms_* ≥ 30). The general trend indicates that the larger a community is, the smaller the relative fraction of experiments needed to get a high accuracy. In fact, similar learning curves could be used as guidelines to determine how many experiments should be performed to reach a desired balanced accuracy.

Variable importance plots are commonly used with random forests to evaluate which variables are the most important to the model by comparing their mean decrease in accuracy scores. Mean decrease in accuracy for each variable is the change in the mean accuracy of the forest predictions when the variable in question is randomly permuted (23). Variable importance is then determined by computing the ratio between their mean change in accuracy and the corresponding standard deviation of accuracy. The randomForest package for R (14) performs this calculation, which we then used to rank exchange reactions as predictors. Variable importance should be interpreted as a ranked list of which variables are generally the most informative rather than a quantification of their effects. These variable rankings are indicators of global predictive importance and can be used to develop insights into why certain features are influential in a general sense. Visualization of the top 20 predictors revealed that the most important predictors tend to belong to the interaction partner (Figure 2F). Interestingly, features from both halves of the interaction vector for the amino acids L-Lysine and L-Glutamate are highly ranked, as are the features for the monosaccharides D-Ribose and Thyminose.

The tree-based approach of random forest to classification can be exploited to determine why specific samples were classified the way they were by examining the feature contributions of each predictor. A feature contribution (Methods) is the quantification of how much a given variable influences the decision of the random forest when a single sample is evaluated. Feature contributions were originally developed for analysis of regression models (24) but have since been adapted for binary classification models (15). In the context of binary classification, feature contributions can be interpreted as how much a feature changes the probability that a given sample is classified as class 1 by the random forest. In the case of our *in silico* data class 1 refers to the non-negative responder type. For our purposes this convention means that a negative feature contribution increases the probability of a sample being classified as a negative responder and a positive feature contribution increases the probability of classified as a non-negative responder. We used the forestFloor package in R (25) to efficiently calculate feature contributions for all 9900 samples.

If the choice of representation faithfully reflects the underlying nature of the system of interest, then feature contributions may be used to gain insight into the underlying mechanisms of an interaction. Were we investigating our *in silico* data as a novel microbial community via pairwise interactions, we would wish to perform additional experiments to identify a metabolite that mediates competition in each negative interaction and then design further detailed experiments to describe molecular details. Absent any particular insight into the nature of the community, metabolites mediating any particular interaction would have to be identified by querying them randomly. In situations where there are few metabolites mediating competition such a process would be a costly endeavor. We wondered if the experimental load could be reduced if for each sample we ranked the 194 metabolites by their net feature contributions and then used the rankings as a guide for the order in which metabolites should be examined (Figure 3A).

**Figure 3.**
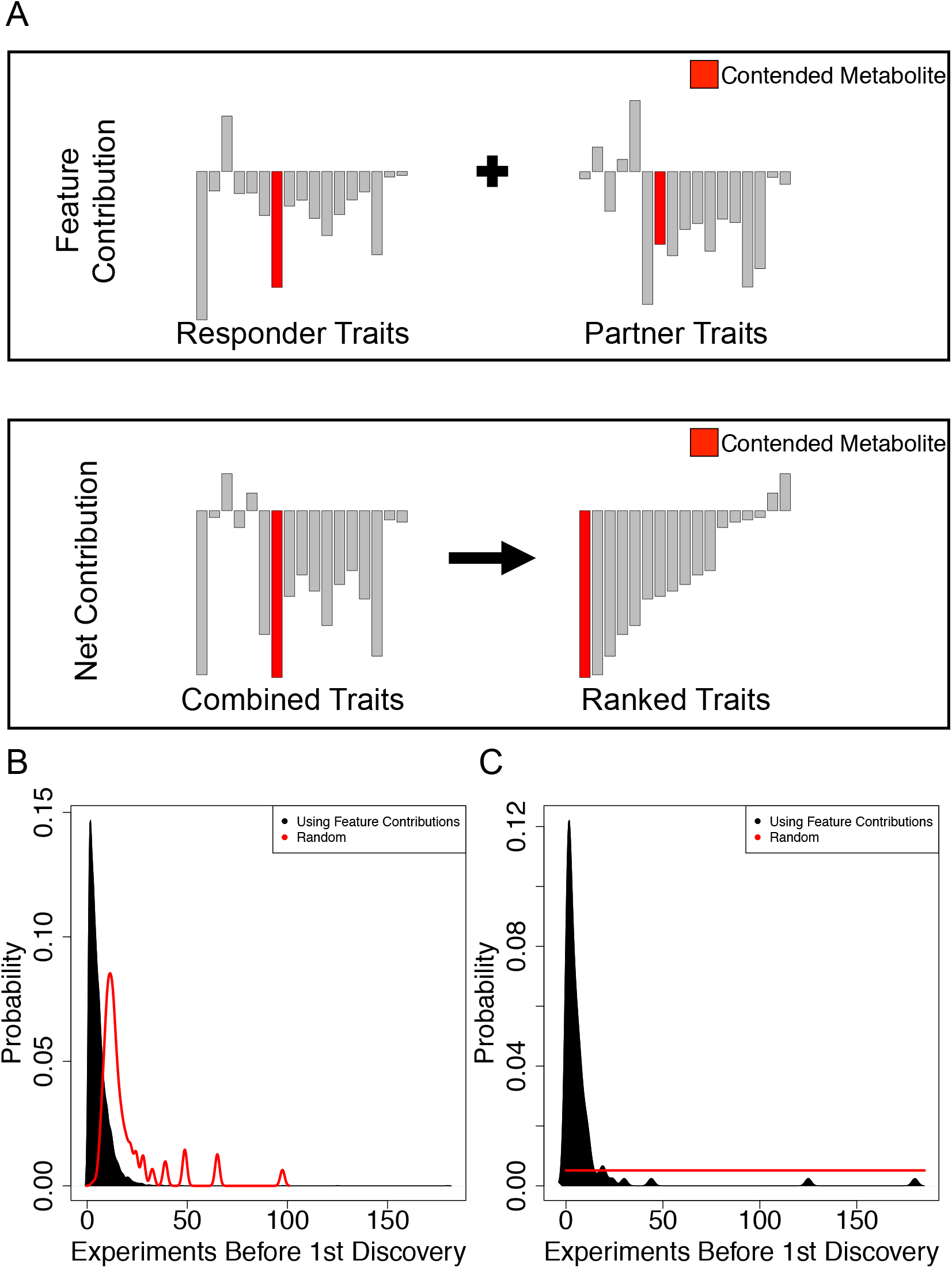
Using feature contributions to find a metabolite for which two organisms compete (also referred to as a contended metabolite) **A.** The first half of the composite trait vector corresponds to metabolite transporters belonging to the organism of interest while the second half corresponds to metabolite transporters belonging to its interaction partner. We were interested in identifying a metabolite that is associated with the negative relative yield for the organism of interest. To establish a ranking of metabolites we took the summation of feature contributions from both halves of the composite trait vector and then sorted the new vector according to the net contribution. Proceeding from the negative end, the rank and identity of the first contended metabolite encountered relative to the negative end of the new vector was recorded. **B.** The probability distribution of the average rank at which the first contended metabolite would be encountered by sampling metabolites randomly one at a time was calculated for each sample and compared to the observed probability distribution. By chance the first metabolite would be encountered after 13 queries. Feature contributions reduce the median number of queries to 4. **C.** 99 samples produced a negative relative through competition for exactly one metabolite. Randomly investigating the 194 candidate metabolites one at a time results in an average of 97.5 experiments before discovering the contended metabolite. Using feature contributions to prioritize the order in which to investigate metabolites instead would typically reveal the contended metabolite on or before the fourth experiment (median = 4).

Because the role of every extracellular metabolite in an interaction is known with COMETS, we were able to evaluate the effectiveness of feature contributions as an experimental guide. For each negative interaction, we identified which metabolites were subject to competition and calculated how many metabolites we would expect to examine at random before encountering any one of the contended metabolites. We then determined the guided rate of discovery by proceeding along the ranked list as determined by net feature contributions and recorded the position at which we first find one of the contended metabolites. Using feature contributions to guide discovery compares favorably to a random selection process. For this community, one would expect on average to perform 19.55 experiments before discovering a relevant metabolite through random selection. Using feature contributions as a guide instead shifted the probability distribution to the left, reducing the expected number of inquiries to 5.54 (Figure 3B). Early discovery of metabolites is particularly valuable when there are very few metabolites mediating an interaction. In particular we wanted to determine how effective feature contributions were in the most challenging cases, those in which a single contended metabolite caused the negative response. We found that for those 99 samples the contended metabolite was found at median rank 4 and in 95 cases was encountered within the top 20 positions (Figure 3C).

Since random forest classifies based on patterns it finds in the data, we can expect that metabolites that tend to be high-ranking predictors are also more likely to be broadly significant to the community than metabolites encountered through random sampling. Across all negative samples there were 109 metabolites that were subjected to competition in at least one interaction. 82 of those metabolites were the first ones encountered using feature contributions, but the majority of encounters were concentrated among eight of them (Supplemental Table 1). DFructose was the most commonly found metabolite, accounting for ~18.7% of all discovered metabolites. Interestingly, fructose has been shown to be implicated in altering the gut microbiome in connection to a number of diseases, including antibiotic treatable (26) metabolic syndrome (27, 28), liver disease (29), and obesity (30).

Notably, our approach is also readily applicable for the discovery of metabolites that mediate positive interactions, which comprise a small minority of all interactions in this specific dataset (420/9900). Due to the scarcity of their occurrence and the dearth of metabolites that mediate positive interactions, discovery of these mechanisms is more challenging. Nevertheless, using ranked feature contributions to find facilitative metabolites was a powerful improvement over a naive approach; requiring a median of 27 queries before discovery versus 65 when selecting metabolites randomly (Supplemental Figure 1).

### Application to a Community of Auxotrophic *Escherichia coli* strains

We analyzed the results of a study in which all-pairwise co-culture experiments for 14 *E. coli* MG1655 auxotrophs were performed (31). In this study, auxotrophic *E. coli* strains were generated by knocking out a single gene that was essential for the production of each of 14 amino acids. Interactions between any given pair of *E. coli* strains are presumably dependent on the direct exchange of the knocked out amino acids, or related precursors (Figure 4A). The total growth of each strain in the 91 experiments was measured after 84 hours and reported as the net fold change relative to the initial inoculum, resulting in 182 total observations. We classified growth phenotypes based on the fold change response a given *E. coli* auxotroph strain had in co-culture with another auxotrophic strain. We used a fold change of 2 as a threshold in order to separate response classes. Fold change 2 is an intuitive threshold since doubling is often used when discussing growth of bacterial populations (32) and classifying all samples with a fold change ≤ 2 as a separate class from those with a fold change greater than 2 results in a balanced data set (Figure 4B). We refer to the 90 growth responses with fold change ≤ 2 as ‘weak’ responses and the 92 growth responses with fold change > 2 as ‘strong’ responses. When discussing a particular *E. coli* interaction, the strain whose response is being predicted is referred to as the ‘receiver’ and its interaction partner is the ‘giver’. We calculated the fraction of interactions that resulted in a weak response for each of the 14 strains and visualized the distribution with a histogram (Figure 4C). Based on this histogram we expect that classification will not be a trivial task because by it appears that the distribution follows either a uniform (Kolmogorov-Smirnov test: p ≈ .93, t test: p ≈ .95) or truncated normal distribution (Kolmogorov-Smirnov test: p ≈ .33, t test: p ≈ .96).

**Figure 4.**
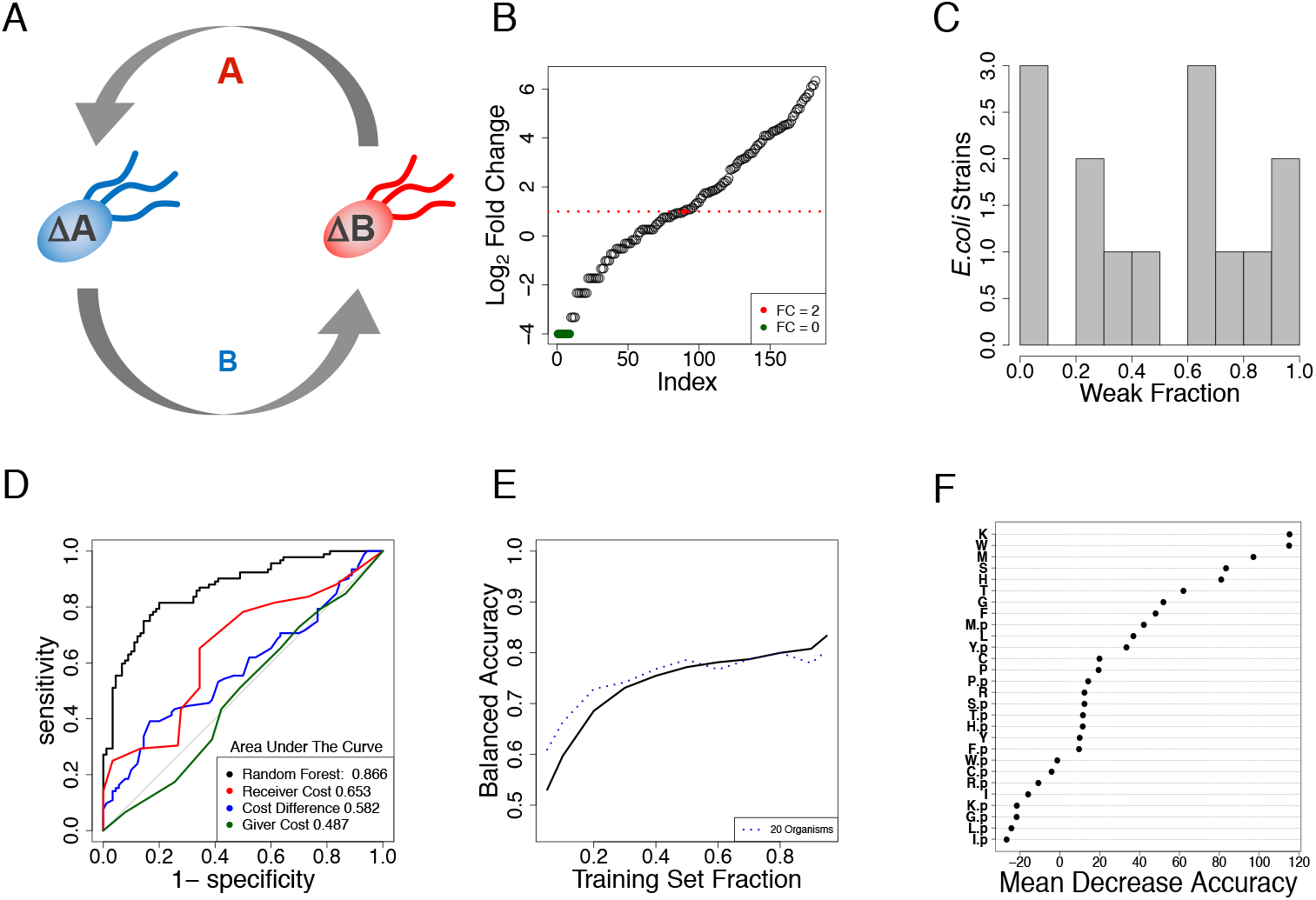
Data representation and results for the case study of a network of auxotrophic *E. coli* strains. **A.** In the original experiment, single gene knockout *E. coli* auxotrophs were co-cultured in a minimal medium. In order for ∆A to grow it must receive amino acid A from ∆B, which in turn must receive another amino acid, B, in order to grow itself. Auxotroph strains were constructed for the following amino acids: cysteine, phenylalanine, glycine, histidine, isoleucine, leucine, methionine, proline, arginine, serine, threonine, tryptophan, and tyrosine. **B.** Sorted fold change for each auxotroph strain across all experiments. *E. coli* strains had a weak response (fold change ≤ 2) 90 times and failed to grow at all 9 times (green circles). In 92 instances the *E. coli* auxotroph population more than doubled over the course of 84 hours. **C.** Histogram of weak responses as a fraction of each strains’ total number of interactions. **D.** ROC curve for random forest classifier using 28 amino acids as predictors for 182 observations. Single value thresholds based on the biosynthetic costs of knocked out amino acids resulted in poorer performance than random forest. **E.** The trajectory of a learning curve built for the *E. coli* interactions closely resembles that of the learning curve for *in silico* communities with 20 organisms. **F.** The 28 amino acids ranked according to their affects on prediction accuracy when randomly permuted. Amino acids corresponding to the receiver strain are enriched near the top of the list. Amino acids are represented by their single letter codes. The suffix ‘.p’ indicates that the predictive feature belongs to the giver strain.

It has been previously observed that biosynthetically costly amino acids tend to promote stronger cooperative interactions than biosynthetically cheap amino acids (31). Given this observation we tested if biosynthetic costs would be useful for predicting whether a strain will have a strong or a weak response in a given interaction. We used the biosynthetic cost of the amino acid provided by the giver strain, needed by the receiver strain, or the difference of these costs as decision thresholds to classify response classes and compared the performance of the random forest classifier to these benchmarks. Since the auxotrophic *E. coli* strains were all derived from the same ancestral strain, only the biosynthetic capabilities for each of the 14 amino acids were relevant predictors for machine learning. As a result, vectors with just 28 elements represented each pairwise *E. coli* interaction. Examination of the corresponding ROC curves shows that biosynthetic costs of amino acids are poor predictors for the qualitative outcome of these *E. coli* auxotroph experiments whereas random forest fares much better as indicated by its ROC curve and by area under the curve (Figure 4D inset). Random forest yielded a balanced accuracy of ~79.2%. Moreover, the trajectory of the learning curve most closely resembles the trajectory of the learning curve for *in silico* communities of 20 members (Figure 4E). Variable importance rankings show that in general, the identity of the receiver’s needed amino acid is often more impactful on classification accuracy than the amino acid that the giver needs, suggesting that specificity of interaction is dominated by auxotrophies, whereas most mutants can in principle provide the missing amino acid (Figure 4F).

Since the pair of knocked-out amino acids determines the interaction for any two auxotrophs, we evaluated how often the corresponding features were the most influential for classification. For each sample, we ranked the feature contributions by the magnitude of their influence and then identified which amino acids were ranked first and second. The feature space is relatively simple and the interaction outcomes are a direct consequence of the relevant auxotrophies. Therefore it is reasonable to expect that the random forest be more strongly influenced by the absence of a predictor than the presence. Of all 182 observations, the knocked out amino acid from the receiver had the largest feature contribution 140 times and the amino acid from the giver was the largest contributor 40 times (Supplemental Table 2). We further found that the top two ranked positions were occupied by the giver and receiver’s amino acids, in any order, 132 of 182 times. Thus the pair of most influential predictors tended to correspond to the underlying mechanism of the interaction, even in instances where the predicted class was incorrect.

Scenarios where the presumed mechanisms are the strongest contributors yet result in misclassification present opportunities to direct research toward interesting outliers in order to understand why they diverge from our expectations. The response of the methionine auxotroph (**Δ**Met) to co-culture with the cysteine auxotroph (**Δ**Cys) was one such case, which is described in detail in Supplemental Figure 2.

## Discussion

Exhaustive pairwise co-culture studies of microbial communities are becoming increasingly common. While such pairwise interactions do not necessarily capture all possible interdependencies in a community (33, 34), they have been shown to be a dominant factor (35), making the reliable prediction and interpretation of predictive models matters of great importance. In this study, we have described a conceptual framework for the representation of microbes and their pairwise interactions in order to address both of these challenges. Our results indicate that representing genome-derived traits of microbes as binary vectors is sufficient for building reliable classifiers for microbial interactions. We also demonstrated two methods for utilizing feature contributions from random forest models to aid in the development of testable hypotheses regarding interaction mechanisms.

Qualitatively predicting the outcome of unobserved interactions is most valuable if those predictions allow precious resources and time to be conserved. To this end the construction of learning curves is an important step in identifying how much data is required in order to achieve desired prediction accuracy from machine learning. Our results indicate that prediction of interactions in communities of all sizes may benefit from machine learning. In situations where the prediction of interaction outcomes is not necessary the properties of random forest makes it a good choice for deciding which traits to study first. Because predictions will always benefit from more data we suggest that if the experimental space is not prohibitively large that all pairwise experiments be performed and feature contributions then be used to derive testable hypotheses. Conversely, when the experimental space is large then one should perform a subset of experiments and incorporate the results of any additional experiments into the training set in order to build new classifiers recursively. The larger the community being studied is the smaller the relative fraction of possible experiments that must be initially performed, these results are promising both for future studies of natural communities, which can contain upward of a thousand unique community members (36) and the assembly of synthetic communities from large libraries of microbes.

Feature contributions provide a clear window into the decision-making process of a random forest when the underlying mechanisms are straightforward, as was the case for the *E. coli* auxotrophs. In cases such as this, interpretation of the model is clear and novel detailed hypotheses can be readily developed. Even incorrect predictions are useful since instances that go against the expectations of the classifier are likely to be worth closer scrutiny. In situations where many factors contribute in a complex manner to an interaction we demonstrated the utility of feature contributions for guiding exploratory experiments by enriching underlying competitive mechanisms near the top of ranked lists. In our particular application of feature contributions to an *in silico* community we used their net effects to identify metabolites that are competed for in negative cases or are used in cross feeding in positive interactions. The particular method of evaluating feature contributions, summing the net effects or sorting them by magnitude is a decision that must be made on a case-by-case basis.

Although we concatenated the binary trait vectors of two organisms in order to form a new composite trait representation, alternative representations of microbes and their interactions should also be explored. Predictions may also benefit from incorporating more information into the base representation of a microbe such as gene copy number or mean transcriptional levels. We also encourage future researchers to acquire information on the same interactions in multiple environmental conditions. Varying the environmental conditions will allow abiotic factors to be incorporated into the representation and should yield additional insight into situations where certain combinations of traits are most relevant.

Going forward we expect that the importance of machine learning in microbial ecology will continue to grow. The need to identify microbial strains that interact in a desired way in any given environment will be one of the most pressing issues for synthetic ecology in the future. Likewise, it would not be surprising if many novel methods for interpreting machine learning algorithms emerge in response to the challenges of understanding the properties of microbial communities.

### Data Availability

Pointers to Datasets obtained from previous work, and used in our analysis are reported in the Materials and Methods Section.

The code necessary to reproduce all our figures and analyses is hosted at: https://github.com/ddimucci/MicrobialCommunities

## Materials and Methods

### Auxotrophic *E. coli* interaction network data

We obtained the measured growth response of individual *E.coli* strains and biosynthetic costs of amino acids from the supplemental files provided by (31).

### Simulation of *in silico* pairwise interactions among gut microbes with dFBA in COMETS

Metabolic reconstructions of human gut associated microbes were obtained from Bauer et al (18). At the time of this writing these models can be downloaded directly from the following URL: https://wwwen.uni.lu/content/download/86230/1056013/file/Bauer_et_al_301_microbe_models.rar

Each metabolic reconstruction encompasses the stoichiometry of virtually all metabolic reactions present in an organism, including uptake/secretion. Flux balance analysis (FBA) is a constraint-based steady state approach that uses this stoichiometry to predict fluxes and growth capacity under a given boundary condition of nutrient availability, and has been described in detail before (19, 37–39). Dynamic flux balance analysis (DFBA) (16) extends classical FBA to perform dynamic simulations in which intracellular metabolites are still assumed to be at steady states, but total biomass and environmental metabolites are treated as time-dependent variables in a discretized approximation. We performed DFBA simulations using our platform for Computation of Microbial Ecosystems in Time and Space (COMETS), which has been previously used to model microbial communities (17). COMETS allows users to implement DFBA in a two-dimensional simulated world populated with multiple metabolic models. We selected 100 metabolic models (18) and identified a common medium that would permit the growth of nearly all models in a monoculture scenario. We then performed all pairwise co-culture simulations of the 100 models using the common media set in a well-mixed batch culture scenario (approximated by using COMETS without spatial structure). For each scenario we saved the record of biomass accumulation and fluxes in order to calculate relative yield and identify mechanisms of interaction, respectively.

### Relative Yield

Relative yield quantifies the change in net growth of an organism in a new environment relative to a reference scenario. For our purpose we compared the growth of models in co-culture to their accumulated growth in monoculture. Relative yield was calculated according to the equation:

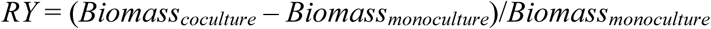

An *RY* < 0 indicates that the responder is detrimentally affected by its partner. Correspondingly, an *RY* of 0 indicates no effect and an *RY* > 0 indicates a positive affect from co-culture with the partner.

### Jaccard Distance

A profile (*model_i_*) encoding the presence (1)/absence (0) of different metabolic reactions in each model *i* was identified based on its stoichiometric matrix. The Jaccard distance (JD) between two metabolic models *i* and *j* was determined with the equation:

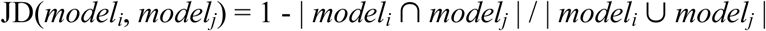

### Representation of interactions with trait-derived features

For a given community *C* the observed co-culture response of each species *i* in the presence of *j* is encoded into the element *X_ij_* of an interaction matrix ***X***. To define a set of trait vectors for each organism in *C*, we start by obtaining a list of *λ* features that can be assigned systematically across all organisms. These features could be the presence/absence of specific genes, functions, or any other trait. In the two case studies reported here, the feature vectors correspond to the presence/absence of specific metabolite transporters in stoichiometric models, and to amino acid auxotrophies, respectively. Thus, each organism *i* is associated with an *λ*-long vector *V_i_*, whose element *k* is 0 or 1 depending on whether or not trait *k* is absent or present in organism *i*. Traits (i.e. elements of the feature vector) that are identical across all organisms are removed from the feature set. Each pair of organisms (*i*,*j*) is then associated with a co-culture feature vector, defined as the concatenation of vectors *V_i_* and *V_j_*, also indicated as *V_i,j_* = [*V_i_*, *V_j_*] (see also Fig. 1). Note that in this concatenated vector, order matters. For the purpose of feeding the data into the random forest algorithm, the complete information about how organism *i* responds in a co-culture with organism *j*, is encoded in the composite feature vector *V_i,j_* and the observed phenotypic response for that specific pair, i.e. *X_ij_*.

### Machine learning

R was used to implement K-nearest neighbor (Knn), Random Forest (RF), and support vector machine (SVM). Knn was implemented with the rknn package, with K set to 3. Library e1071 was used for SVM and we used a linear kernel and cost=0.1. Random forest was implemented with the randomForest library. All three algorithms had similar performance on the *E. coli* data set but RF was significantly better on the *in silico* data set (data not shown).

Random forests are ensemble classifiers that aggregate the results of many individual decision trees. Each tree in a random forest is assigned a synthetic data set that is of the same size as the training set but generated through sampling with replacement. The result is that the average tree is trained on approximately 2/3 of the observations; these observations are referred to as in bag samples. The remaining 1/3 of observations not in the synthetic data sets are referred to as out of bag samples. The new synthetic data set is placed at the root node of a new tree; next a randomly selected subset of predictive features is queried for the best split of the data into two child nodes. This process is repeated at each node until a stopping criterion is met. Classification accuracy of individual trees is assessed by running their out of bag samples down the tree and recording the results. See (19) for a full description of the algorithm.

### Evaluation of classifiers

We used balanced accuracy to evaluate performance of classifiers. This metric is based on the values from the confusion matrix: true positive (TP), true negative (TN), false positive (FP), and false negative (FN). Balanced accuracy is calculated as follows:

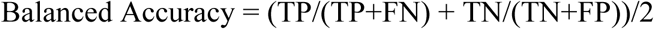

### Learning curves

To construct learning curves we defined a set of fractions, *fr* = [.05, .1, .2, .3, .4, .5, .6, .7, .8, .9, .95] where we would evaluate balanced accuracy of the model using cross validation. For all cross-validation experiments we ensured that observations X_ij_ and X_ji_ were either both in the training set or in the test set. For each fraction in *fr* we randomly selected max(1/*fr_i_*, 1/(1-*fr_i_*)) mutually exclusive subsets of the data to use as a training set or as a hold out test set if *fr_i_* > .5. This process was repeated until at least 10 subsets of the data were selected for each fraction. The median balanced accuracy of the classifier was then calculated at each point along *fr.* In order to investigate the effect of the community size on the learning curve we defined a set of community sizes *S* = [10, 20, 30, 40, 50, 60, 70, 80, 90]. For each community size *S_i_* we randomly selected five community sub-matrices, *X_k_*, from the full *in silico* community matrix (*X*). Then for each *X_k_* a learning curve was determined and the median learning curve for balanced accuracy of each community size was calculated

### Feature contributions for binary classifications

Each tree in the random forest is presented with a bootstrapped subset of the provided data as a training set that is used to grow the tree. The training set is placed at the root node of the tree. The fraction of instances in the training set of class *C_1_* is 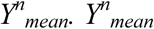 is the probability that an instance selected at random from the root node is assigned to class *C_1_*.

There are two steps in the calculation of feature contributions for a new sample: First calculate the sum of all local increments in each tree, second determine the average contribution over the forest. A local increment for a feature *f* is calculated if the split was performed on *f* and is defined as the change in probability of *C_1_* in a parent node (p) and a child node (c):

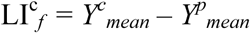

Local increments quantify the change in the probability of a sample being classified as *C_1_* between the child node and the parent node when *f* is the splitting criterion. The feature contribution, 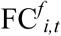 of feature *f* in tree *t* for observation *i* is the sum of all LI_f_ along the path from the root node to the terminal node. 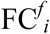 is given by the equation:

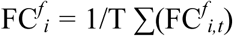

Feature contributions were calculated with the forestFloor package and are calculated using only trees for which a sample was out of bag.

### Application to positive samples

Positive samples constituted a small minority of the total samples (420/9900) and result in a heavily imbalanced data set when we try to classify these samples as separate from either neutral or negative responses. Imbalanced datasets often result in very high classification accuracies because the classifier simply predicts every instance to be of the majority class. This is an issue because if we are to have confidence in feature contributions an adequately performing classifier is desired. In order to develop a reliable classifier we under sampled the non-positive instances by randomly selecting a subset of 420 of them to pair with the 420 positive observations and then trained a random forest on the new data set. To estimate the efficacy of the random forest we repeated this process 100 times and note that there was a median balanced accuracy of ≈ 85%. We then used a single random forest model to calculate feature contributions in the context of identifying positive responses.

## Acknowledgments

We are grateful to Dr. Jenny Bhatnagar and Dr. Carolyn Zeiner whose work with fungal interactions served as the initial inspiration for this study and to members of the Segrè lab for helpful discussions and for feedback on the manuscript. DS and DD acknowledge funding from the Defense Advanced Research Projects Agency (Purchase Request No. HR0011515303, Contract No. HR0011-15-C-0091), the U.S. Department of Energy (DE-SC0012627), the NIH (T32GM100842, 5R01DE024468, R01GM121950 and Sub_P30DK036836_P&F), the National Science Foundation (1457695 and NSFOCE-BSF grant 1635070), the Human Frontiers Science Program (RGP0020/2016), and the Boston University Interdisciplinary Biomedical Research Office.

