## Supplementary Materials for "Machine learning reveals missing edges and putative interaction mechanisms in microbial ecosystem networks" (from another version of this preprint: version 1, 2018-03-21)

**Supplemental Table 1**

Mechanistic metabolites found first in negative interactions using ranked feature contributions as a guide.

| Name | Times Found First | Median Rank |
|---|---|---|
| D-Fructose | 1041 | 4 |
| gly-asn-L | 347 | 9 |
| ala-L-asp-L | 308 | 8 |
| gly-asp-L | 261 | 8 |
| L-Glutamine | 252 | 3 |
| L-Lysine | 245 | 2 |
| Deoxyguanosine | 197 | 2 |
| Inosine | 192 | 3 |
| G3P | 176 | 2 |
| Deoxycytodine | 161 | 3 |
| Maltose | 151 | 6 |
| Aminoethanol | 149 | 2 |
| D-Ribose | 145 | 2 |
| Formate | 142 | 2 |
| L-Proline | 116 | 6.5 |
| D-Glucose | 113 | 5 |
| Spermidine | 99 | 5 |
| Deoxyinosine | 98 | 3 |
| L-alanglycine | 86 | 6 |
| Riboflavin | 82 | 5 |
| L-Glutamate | 80 | 4 |
| meso-2,6-Diaminopimelate | 75 | 1 |
| Adenosine | 66 | 3 |
| Cytidine | 66 | 4 |
| L-Valine | 59 | 4 |
| N-Acetyl-D-glucosamine | 58 | 3 |
| Fumarate | 58 | 1 |
| Cys-Gly | 51 | 3 |
| Gly-Tyr | 47 | 11 |

| | | |
|---|---|---|
| Gly-Phe | 47 | 12 |
| L-Isoleucine | 44 | 5 |
| ocdca | 33 | 2 |
| L-Arginine | 32 | 2 |
| ala-L-glu-L | 29 | 9 |
| gly-pro-L | 28 | 11.5 |
| Deoxyadenosine | 26 | 6 |
| L-Lactate | 24 | 1 |
| Gly-Leu | 23 | 10 |
| Uracil | 20 | 2.5 |
| PAN | 20 | 6 |
| Galactose | 19 | 3 |
| Guanosine | 18 | 1 |
| Uridine | 17 | 2 |
| L-Alanine | 17 | 2 |
| Putrescine | 17 | 3 |
| Gly-Met | 16 | 2 |
| L-Serine | 15 | 4 |
| TRHL | 14 | 2 |
| XAN | 13 | 6 |
| Ala-Leu | 13 | 8 |
| Thymidine | 13 | 4 |
| ala-L-Thr-L | 11 | 9 |
| NH3 | 11 | 7 |
| Citrate | 10 | 7.5 |
| Succinate | 10 | 1 |
| Ornithine | 9 | 12 |
| D-Alanine | 8 | 4 |
| Ala-His | 8 | 9.5 |
| Menaquinone 7 | 8 | 4.5 |
| L-Methionine | 6 | 6.5 |
| Folate | 6 | 3 |
| LACT | 5 | 3 |
| HYXN | 5 | 10 |
| Nicotinamide ribonucleotide | 5 | 8 |
| Nicotinamide | 5 | 3 |
| Heme | 5 | 1 |
| Thiamin | 5 | 3 |

| | | |
|---|---|---|
| L-Tryptophan | 4 | 7 |
| Sucrose | 3 | 4 |
| L-Leucine | 3 | 7 |
| H2S2O3 | 2 | 2.5 |
| CELB | 2 | 14.5 |
| L-Phenylalanine | 2 | 2.5 |
| Dephospho-CoA | 2 | 15.5 |
| GLUM | 2 | 5.5 |
| L-Threonine | 1 | 1 |
| Isoprene | 1 | 2 |
| met-L-ala-L | 1 | 98 |
| 4-Hydroxybenzoate | 1 | 3 |
| D-Arabinose | 1 | 2 |
| 2-Oxoglutarate | 1 | 1 |
| L-Malate | 1 | 12 |

**Supplemental Table 2**

Position of receiver and giver knocked-out amino acids when ranked according to the magnitude of their feature contributions.

| **Predictive Rank of Knocked Out (KO) Amino Acids** | **Total Occurrences (Fraction of samples)** |
|---|---|
| Receiver KO 1st | 140 (.769) |
| Giver KO 1st | 40 (.22) |
| Receiver KO 2nd | 35 (.192) |
| Giver KO 2nd | 97 (.533) |
| Receiver KO 1st/Giver KO 2nd | 97 (.533) |
| Giver KO 1st/Receiver KO 2nd | 35 (.192) |

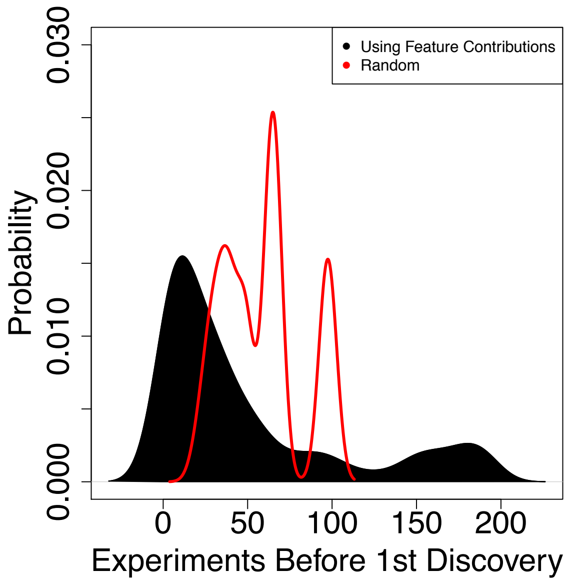


**Supplemental Figure 1.** Using feature contributions to find a facilitative metabolite. The probability distribution of the average rank at which the first facilitative metabolite would be encountered by sampling metabolites randomly one at a time was calculated for each sample and compared to the observed probability distribution obtained from using ranked feature contributions. By chance the median first metabolite would be encountered after 65 queries (mean ≈ 58.7). With feature contributions the median number of queries was 27 (mean ≈ 50).

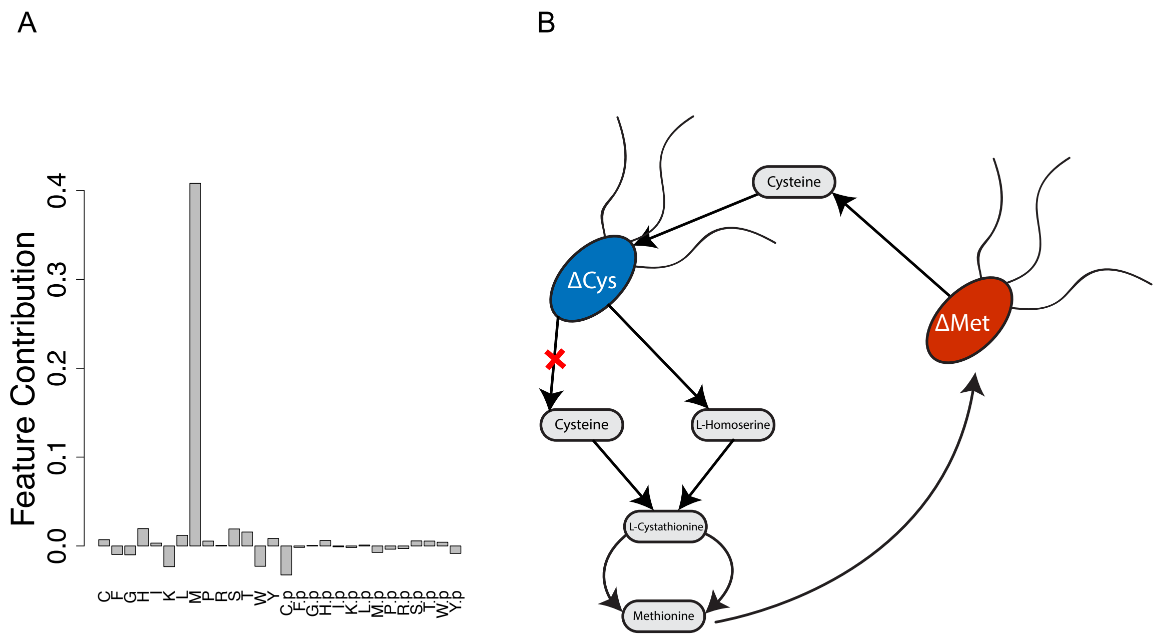


**Supplemental Fig. 2. A.** Bar plot of the calculated feature contributions for the growth response of ΔMet co-cultured with ΔCys, in the *E. coli* auxotrophs case study. The feature contribution from the receiver methionine is ≈ .41. The feature contribution from the giver cysteine is ≈ -.03. The net contribution from the remaining predictors is ≈ .02. **B.** ΔMet typically had a strong response in co-culture no matter the identity of its interaction partner; 12 interactions resulted in strong response type for ΔMet. When it was grown with ΔCys, however, it had a weak growth response. The use of feature contributions correctly identified the receiver's methionine and the giver's cysteine as the first and second most important predictors, respectively, in this interaction. The contribution from the receiver methionine is overwhelmingly positive; reflecting the fact that ΔMet typically benefits strongly from co-culture and results in a strong response prediction (A). In order to develop a hypothesis for why this interaction defied the expectations of the random forest we consulted the literature regarding the biosynthetic pathways for methionine and cysteine. We learned that under the specified growth conditions cysteine is necessary for the biosynthesis cytathionine. Cytathionine is subsequently required for the biosynthesis of homocysteine, which is in turn required for the production of methionine. Given this knowledge we suspect that **Δ**Cys is unable produce methionine until enough cysteine accumulates in the environment and **Δ**Met must wait for ΔCys to use **Δ**Met's own excess cysteine to produce excess methionine (B). ΔMet is unlikely to be able to provide an abundance extracellular cysteine for ΔCys early in the interaction because cysteine biosynthesis is tightly regulated due to its toxicity (36) but efflux of cysteine has been proposed as a potential regulatory mechanism (37) and likely allows ΔMet to produce low levels of extracellular cysteine. The waiting time associated with ΔMet derived cysteine accumulating in the environment would result in a delayed start to growth for both strains relative to other interactions involving ΔMet. The reported fold changes in this experiment were fold 0.8 for ΔMet and 1.2 for ΔCys. The weak growth response of ΔMet is consistent with the idea of a delayed start to growth.
